# Combined morphological and transcriptomic analyses reveal genetic regulation underlying the species-specific bulbil outgrowth in *Dioscorea alata* L

**DOI:** 10.1101/585364

**Authors:** Zhi-Gang Wu, Wu Jiang, Zheng-Ming Tao, Xiu-Zhu Guo, Xiao-Jun Pan, Wen-Hui Yu

**Affiliations:** Key Laboratory for Plant Genetic Improvement, Institute of Subtropical Crops, Zhejiang Academy of Agricultural Sciences, Wenzhou 325005, China; School of pharmacy, Wenzhou Medical University, Wenzhou 325035, China; Quzhou Academy of Agricultural Sciences, Quzhou 324000, China

**Keywords:** Bulbil, genetic regulation, outgrowth, transcriptome, yam (*Dioscorea alata* L.), phytohormone signals

## Abstract

In yam (*Dioscorea* spp) species, bulbil at leaf axils is the most striking species-specific axillary structure and exhibits important ecological niches as well as crop yields. Genetic regulation underlying bulbil outgrowth remain largely unclear. We here first characterized the development of bulbil from *Dioscorea* alata L. using histological analysis and further performed full transcriptional profiling on its key developmental stages. Comprehensive mRNA analyses suggested that long-distance phytohormone signals including auxin, CK and ABA, play critical roles in controlling the initiation of bulbil through coordinately altering expression levels of genes involved in localized hormone metabolism and transport. Sucrose functioned as a novel signal and was required strongly at the early stage of bulbil formation, thus promoting its outgrowth through up-regulating trehalose-6-phophate pathway. GO pathway analysis demonstrated that genes are enriched in biological processes related to light stimuli, cell division, cell wall modification and carbohydrate metabolism. Particularly, some novel genes including dioscorin A/B, starch synthetic enzymes and chitinases showed remarkably high expression levels and strengthened the outgrowth of bulbil. Our data set demonstrated that the initiation of bulbil was highly regulated by a large number of transcriptional regulators. RNA in situ hybridization with MYB, WRKY and NAC transcription factors confirmed their key roles in triggering bulbil initiation. Together, our findings provide a crucial angle for genetic regulation of controlling the unique reproductive development of bulbils. Transcriptome data set can serve as a valuable genomic resource for yam research community or further genetic manipulation to improve bulbil yields.

**Highlight:** Transcriptomic data identified multiple functional genes and regulators; long-distance signals (auxin, CK, ABA), and sucrose as a novel signal play critical roles in controlling bulbil outgrowth.

## Introduction

Unlike animals, plants are sessile and evolve a high degree of plasticity in architectural forms to respond to environmental changes (Barthélémy and Caraglio, 2007). Such developmental plasticity is mainly determined by the shoot branches, which grow from buds, arising the axillary meristems (AMs) at the axil of leaf primordia (Evers *et al*., 2011). Axillary branching is genetically regulated, and generally modified by biotic and abiotic factors to establish a unique form suitable for the ecological/evolutionary niches in which the plant grows. In many species, branches or other species-specific axillary structures are key components of agricultural traits that control the plant biomass and crop yields (Li *et al*., 2003; Evers *et al*., 2011).

Over the past 30 years, most of our understanding in molecular regulatory pathways for plant branching establishment, is derived from model species such as *Arabidopsis thaliana*, rice and maize, or from pea (*Pisum sativum*) and rose (*Rosa hybrida*) plants (Gallavotti *et al*., 2010; Domagalska and Leyser, 2011). Auxin has been considered as a central signal in controlling the initiation of axillary meristem (AM) and the regulation of secondary branching (De Smet and Jürgens, 2007). Two widely accepted hypotheses, auxin transport canalization (ATC) and the second messenger (SM) models, have been proposed to describe the role of auxin in regulating the bud development (Prusinkiewicz, *et al*., 2009). Central to ATC model is that auxin from shoot apex inhibits the outgrowth of buds, and must be depleted to activate new AM expansion, in which the establishment of a polar auxin transport (PAT) stream is essential and driven by a positive expression of PIN efflux proteins (Balla *et al*., 2011). SM model contend that auxin does not enter the bud, and regulates the production of branches by second messengers, which moves directly into the bud to control its activity (Domagalska and Leyser, 2011). As a good candidate for second messenger, cytokinin (CK) synthesized in the roots that is regulated negatively by the auxin flow in the main stem, travels to the shoot and enters the bud where it promotes bud outgrowth (Ferguson and Beveridge, 2009). Important studies revealed that CK enables buds to escape dormancy by a direct application in many plants, even in the presence of apical auxin (Elfving and Visser, 2007; Roman *et al*., 2016). Instead, several reports indicated that abscisic acid (ABA) has a strong negative correlation with the rate of bud outgrowth, especially in later developmental stages (Reddy *et al*., 2013; Yao and Scott, 2015). The ABA-induced inhibition is verified by exogenous ABA application in decapitated plants (Cline and Oh, 2006). Transgenic experiments disclosed that ABA-insensitive mutants (*abi*1, *abi*2, *abi*3) in Arabidopsis enhance the outgrowth of lateral buds (Zheng *et al*., 2015).

More recently, there is a large body of physiological evidences from many species, supporting sugar (especially sucrose) as a novel signal required for the triggering of bud outgrowth (Evers, 2015). Exogenously supplying sucrose in pea plant stimulates the outgrowth of buds by suppressing the expression of BRANCHED1 (BRC1) inhibitor gene in buds (Mason *et al*., 2014). The sugar-mediated transduction mechanism has been proposed to be linked to influence the biosynthesis, transport of certain hormones, such as auxin, CK, and SL, but remain to be further discovered (Barbier *et al*., 2015). The plasticity in branches is also highly sensitive to environmental inputs, especially to light conditions (Kebrom and Mullet, 2015). An exposure to a low light intensity or a low far-red (R/FR) ratio light inhibits the bud outgrowth, even when buds are freed from apical dominance (Holalu and Finlayson, 2017). This suggests that light is essential and acts as a morphogenic signal in triggering bud organogenesis (Roman *et al*., 2016). The phenomenon is well characterized in plant shade avoidance syndrome (SAS), where phytochromes (especially phy B) perceive and monitor the low R/FR light, thereby decreasing bud numbers (Reddy *et al*., 2013). Important evidences have proved that phy B can transduce largely the effects into changes in expression of genes associated with hormone biosynthesis and signaling, or bud growth (Reddy and Finlayson, 2014; Kebrom and Mullet, 2016).

Despite the considerable advances that have accumulated over many years in model plants, the full regulatory mechanism remains to be explored to understand the complexity of branching patterns. Especially for the species-specific axillary organs, the regulatory pathways is distinct between different species (Evers *et al*., 2011). The genus *Dioscorea* has been considered to be among the most primitive of the Angiosperms and differentiated species (Burkill, 1960). In more than 600 members recorded in this genus, the most striking branch-plasticity is the occurrence of bulbils that are termed as minor storage organs or aerial branches (aerial tubers), and initiate at their leaf axils (Wickham *et al*., 1982; Murty, 1983). Ecologically, this structure enables the plants to spread rapidly and engulf native vegetation (Mizuki and Takahash, 2009). In practice, yam bulbil serves as an effective means for vegetative reproduction, and are often dormant and will germinate when drop to the ground in the following season (Main *et al*, 2006). In many yam species, aerial bulbils are most common and have been produced widely as foods or pharmaceutical uses (Fu *et al*., 2006). Despite its importance in commercial values and ecological plasticity, the gene regulatory network throughout bulbil development remain largely unclear. Here, we selected a typical yam species *(Dioscorea alata* Linn.) and performed full transcriptional profiling on key developmental stages during bulbil outgrowth by combining morphological analysis. The expression of genes involved in the control of bulbil outgrowth, including hormone-, sugar-, light-, and cycle-related genes and other novel genes, was identified to decipher the regulatory pathway underlying the species-specific bulbil outgrowth. Together, the genome-scale gene expression profiling investigated provide valuable genomic resources for further biochemical discovery and functional analysis for yam bulbil development.

## Materials and methods

### Plant materials

The homogenetic seedlings of yam (*Dioscorea alata* L.) were asexually propagated using severed tubers (100∼150 g per plant) from an identical mother plant. The seedlings were grown in Wenzhou botanical garden (Wenzhou city, Zhejiang province, China) under standard conditions. The bulbils were seen in the axil of leaf primordia after 130 days growth. Four staged samples of bulbil were harvested at 0 day after bulbil emergence (0 DAE, labeled T1), early stage (7 DAE, T2), middle stage (15 DAE, T3), mature stage (35 DAE, T4). Five plants were collected for each bulbil RNA sample, and then immediately frozen in liquid nitrogen for Illumina RNA-seq.

### Histological analyses of bulbil development

Bulbils across the four investigated stages were fixed in FAA solution (70% ethanol: formalin: acetic acid= 90:5:5) for 24 h at 4 °C. The fixed bulbils were dehydrated through a graded ethanol series, and embedded in paraffin, as described previously (Xing *et al*., 2010). Longitudinal sections of 10 μm were prepared with a rotary microtome and stained in safranin-alcian green according to standard histological procedures (Gutmann,1995). Stained sections were observed and documented under an OLYMPUS BX60 light microscope (Olympus, Japan).

### Determination of sucrose content

Bulibls were ground in liquid nitrogen. Sucrose was extracted with 80% ethanol for 40 min at 60 °C in cap-sealed tubes using 1 mL per 200 mg powder sample. The extraction was carried out twice with the same condition. The combined suspension was centrifuged at 10,000 *g* for 10 min at 4 °C. The supernatant was analyzed and sucrose contents were determined with HPLC-ELSD analysis (Agilent 1200), as described previously (Ma *et al*., 2014). Five independent experiments were carried out for each staged bulbils.

### RNA isolation and sequencing

Total RNA of each bulbil was isolated using TRIzol reagent (Invitrogen) and purified using DNase1 (TURBO DNase, Ambion, USA). The integrity of RNA (RIN>8.5) was detected using a Bioanalyzer 2100 (Agilent Technologies, Santa Clara, CA). RNA-seq libraries were prepared using the cDNA Synthesis kit (Illumina Inc., San Digo, USA) following the standard Illumina preparation protocol. Paired-end sequencing (2×150 bp) was conducted with Illumina HiSeq 2500 (Illumina Inc., San Diego, USA) by Biomarker Biotechnology Corporation (Beijing, China). Three independent biological replicates were analyzed for each staged samples.

### Transcript assembly and bioinformatics analysis

The raw reads were refined by removing reads with only adaptor and unknown nucleotides>10%, and low-quality reads with average Phred quality score<30. The clean reads were used for robust de novo assembly of a set of transcriptome using Trinity software package (version R2013-02-25) (Haas *et al*., 2013). Subsequently, we estimated expression abundance of each transcript by calculating FPKM value (Fragments per kilobase of exon per million fragments mapped) using TopHat (version 2.0.8) (Trapnell *et al*., 2012), and those with FPKM >1.5 were remained for further analysis. Pearson correlation coefficient of expression levels between three biological replicates was calculated to assess the reliability of sample collection using cor R package.

Differentially expressed genes (DEGs) between differentially staged bulbils were identified using Edge statistical test in terms of the following criteria: false discovery rate (FDR) <0.01, and an absolute expression fold-change≥2 for a given genes between any two staged samples (Robinson *et al*., 2010). We annotated biological function for DEGs using NR public databases according to BLASTX analysis with a cut-off E-value of 10^−5^. GO slim was conducted to obtain GO annotations using Blast2GO (Conesa *et al*., 2005). To obtain enriched slims, we further performed GO term enrichment analysis using the algorithm and Kolmogorov-Smirnov (KS) test (P -value≤0.001) in R package topGO (Alexa *et al*., 2006). Based on all expressed genes (FPKM >1.5), principal component analysis (PCA) was carried out to explain the relatedness among all staged samples. According to certain functionally categorical genes, we performed hierarchical cluster analysis (HCA) to present gene expression pattern using pheatmap library in R software. In addition, transcription factors (TFs) were identified in terms of Arabidopsis transcription factor database (Perez-Rodriguez *et al*., 2010). A P parameter was defined as the expression proportion of certain TF family, and calculated as described previously (Yu *et al*., 2015).

### Validation of Genes Using Quantitative RT-PCR

Quantitative reverse transcription-PCR (qRT-PCR) was performed to verify transcriptomic profiling results with twenty selected genes. Three biological replicates were conducted for each gene. The PCR amplification primers for selected genes were designed with Primer3 software, and sequences were listed in Supplementary Table S1. RNA was extracted and treated as described above. The cDNA was prepared with SuperScript III Reverse transcriptase (Invitrogen) following the manufacturer’s protocol on 1μg of RNA. The qRT-PCR amplification was run in an ABI 7500 HT (Life technology) with SYBR Green I Master Mix (TaKaRa), and the reaction mixture and program was carried out as described in our previous work (Wu *et al*., 2014). Quantification of gene expression was normalized using EF-1a gene (Accession: JF825419) as an internal control, and counted according to the 2^− ΔΔCt^ method (Livak and Schmittgen, 2001).

### RNA in Situ Hybridization

RNA in situ hybridization was performed using the method described by Siciliano *et al*. (2007). Each staged bulbils were fix in FAA and embedded in paraffin as described above. Gene-specific fragments for probe synthesis were amplified by PCR using designed primers: 5’-GAAGAGCACCATGCTGTGAG-3’ and 5’-TAATACGACTCACT ATAGGGCCACATCTCAGCAATCCAG-3’ for MYB (Gene ID: c126446.graph-c0), 5’-TGGAGAGCCTTTGATCGGTT-3’ and 5’-TAATACGACTCACTATAGGGCCAC TGCTCTAAACGAAGG-3’ for WRKY (c125026.graph_c1), and 5’-AGTGCAT TACCTCTGCCGGA-3’ and 5’-TAATACGACTCACTATAGGTACCTGGCA ATTCCCAAGGA-3’ for NAC (c116834.graph_c0). The resulting PCR fragments were used as templates for synthesis of digoxigenin (DIG)-labeled antisense and sense riboprobes with the T7/SP6 riboprobe and a DIG-RNA labeling mix (Roche). Sections (8-10 μm) were treated with 1 μg mL^−1^ proteinase K for 30 min at 37°C, and then washed under stringent conditions as described previously (Hsu *et al*., 2015).

## Results

### Morphology of bulbil during its developmental stages

Yam bulbils are generated naturally on the leaf axil when the main apex stops growing (Fig. 1A). To characterize the developmental process of bulbils in detail, we designed four growing sequences based on our observation and important publication (Murty and Purnima, 1983), including the initiation (T1), early (T2), middle (T3) and mature (T4) stages of bulbil formation (Fig. 1B). At T1 stage, the cells of 2-3 layers below the leaf axil have undergone periclinal and anticlinal divisions, and further developed into a hump-like meristematic tissue that is termed as the bulbil primordium (BP). The BP at this stage was pivotal for subsequent bulbil outgrowth and still remained differentiated in part. Meanwhile, a dome-shaped organ was visible in the leaf axil. At T2 stage, the cells from BP meristematic zone (Fig. 1H) became highly meristematic and showed successive cell division and enlargement, further formed young bulbil. The young bulbil was top-shaped and smooth on the surface, and the root primordia (RP) was seen in cortical region at this stage (Fig. 1G). At the next stage (T3), the size of bulbil was enlarged rapidly, as the meristem in the central region of bulbil is continuously widen. Increased cell numbers and enlarged volume by filling starch grains shaped quasi-round bulbils. The bulbil at this stage had a distinct peripheral region covered by several rough epiderm layers (Fig. 1B). At T4 stage, the activity of meristematic tissue was depleted mostly and mature bulbils reached 1.2-3.0 cm in diameter. Multiple RPs were emerged on bulbil surface, which enable bulbils to spread rapidly in next season. Together, bulbil morphology at the four developmental stages was distinct.

**Fig. 1.**
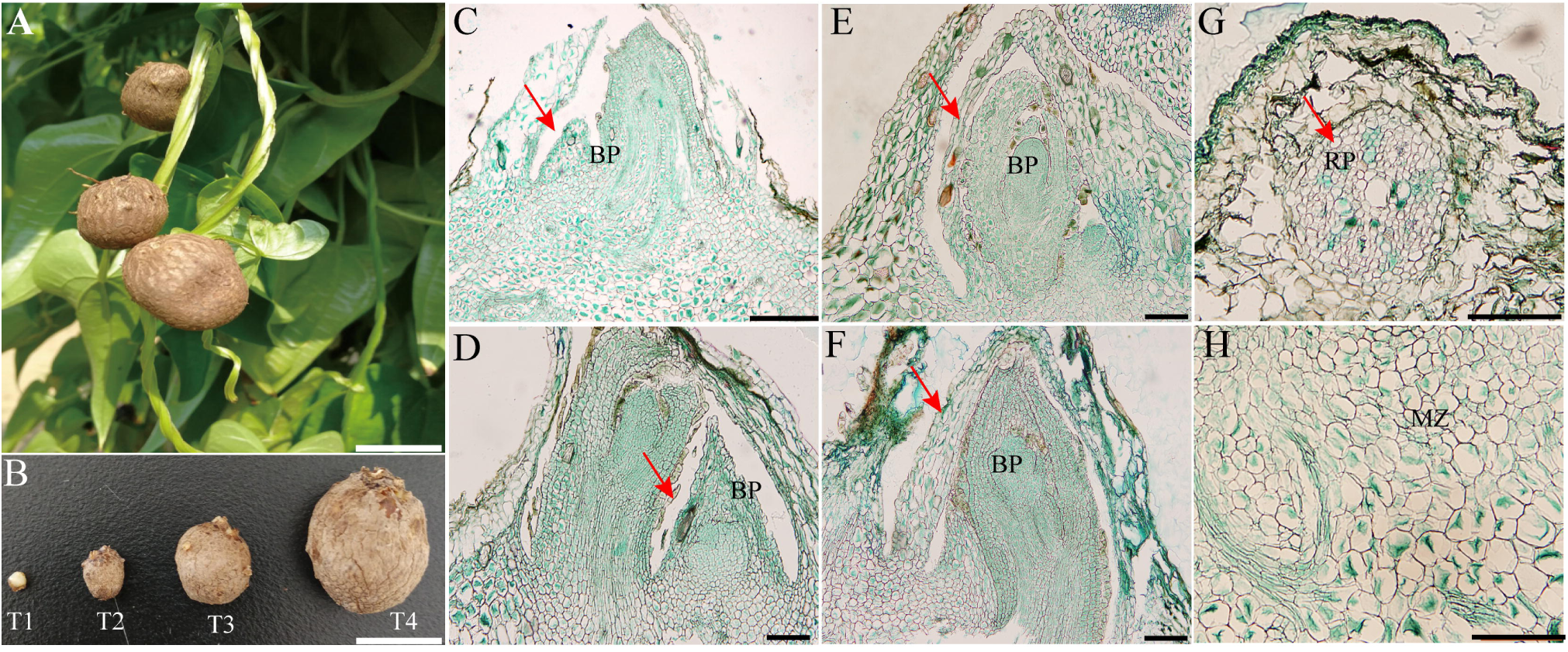
Morphology of bulbil during key developmental Stages. **(A)** Bulbil phenotype. **(B)** Photographs of bulbil at the initiation (T1), early (T2), middle (T3) and mature stages (T4). **(C-F)** Paraffin sections of bulbils at T1 (C), T2 (D), T3 (E), T4 (F) stages. The images show the zone of the junction region between bulbil and axil. G, Showing root primordia (RP). D, Showing meristematic zone (MZ). Bras: A and B, 1 cm; C, 500 µm; D to H, 200 µm.

### Transcriptome profiling during bulbil outgrowth

We sequenced 12 RNA libraries from bulibls at its key developmental stages (T1,T2,T3,T4) based on observation above. All raw reads obtained have been submitted to NCBI under accession number SRP152752. After stringent data cleanup and quality checks, a high quality of RNA-Seq data was obtained for each sample with a quantity of 49 million to 66 million paired-end reads (Supplementary Table S2). The de novo assembly generated 199,270 transcripts, approximately 36% of transcripts were in the size range 200-500 bp (Supplementary Table S3). The homologous transcripts were further clustered with > 95% similarity, the final bulbil transcripts generated 97,956 unigenes with a total length of 79,527,036 bp and an average length of 812 bp.

Based on whole gene expression profile (FPKM values), PCA visualized four stage-specific clusters with the first two components explaining 77.4% of total variance, and revealed distinct mRNA populations between different staged bulbils (Fig. 2A). We identified that 752 (in T1), 659 (T2), 385 (T3) and 1210 (T4) genes that showed highly stage-specific expression patterns, but most genes were shared in all staged bulbils (Fig. 2B). Meanwhile, we assessed gene expression profiles between biological triplicates and they were highly correlated (*R*^*2*^> 0.83), indicating that each staged bulbils were well collected (Supplementary Fig. S1). To confirm the reliability of RNA-seq, we further performed a more rigorous expression measure for twenty selected genes using qRT-PCR analysis. We disclosed a good agreement with a high linear correlation (*R*^*2*^>0.80; Supplementary Fig. S2) between RNA-seq and qRT-PCR technologies.

**Fig. 2.**
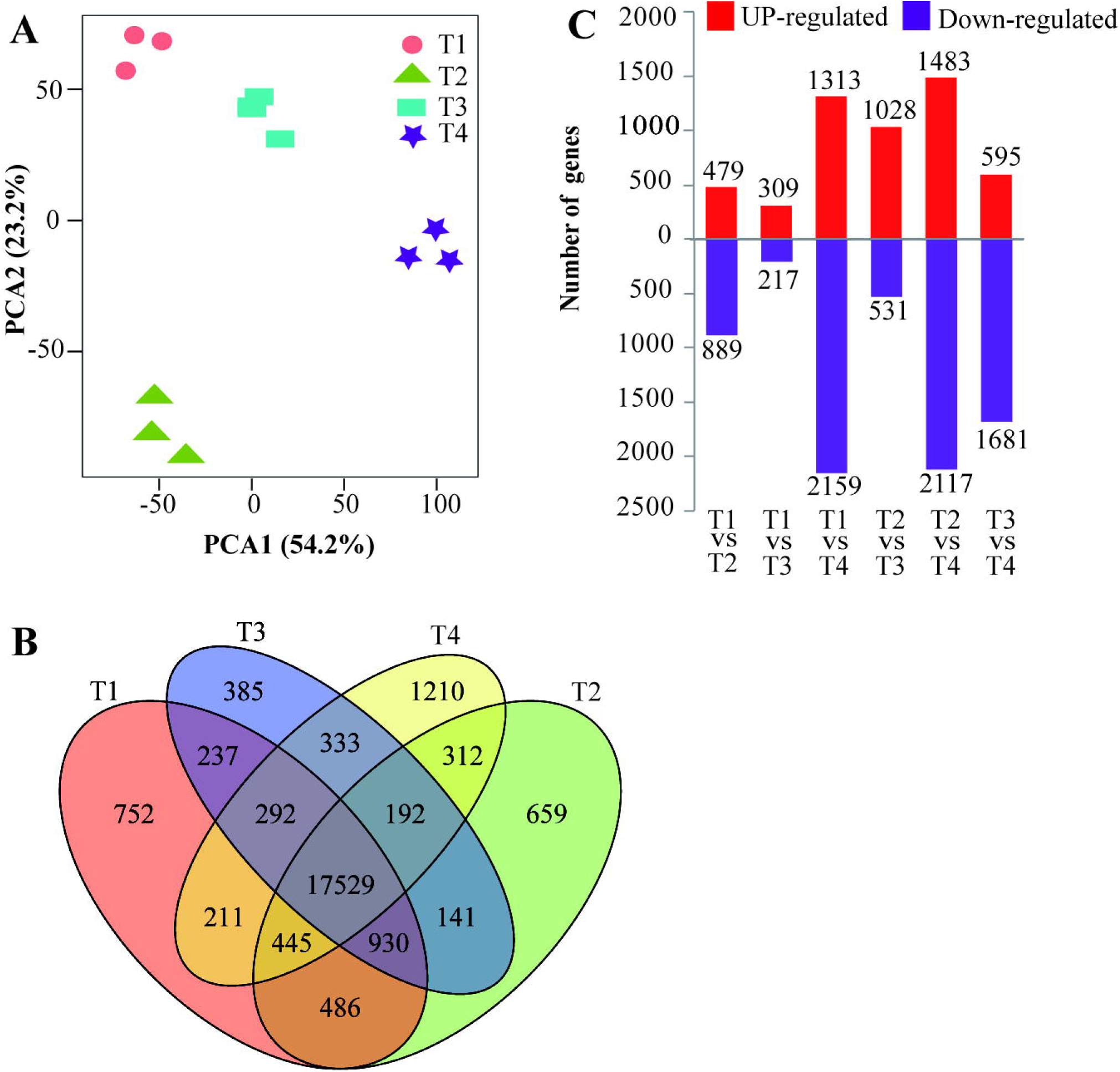
Gene expression profiles in developing yam bulbils. **(A)** Principal component analysis for 12 bulbil samples shows four stage-specific groups based on all gene expression data. **(B)** Venn diagram shows the numbers of unique and overlapping expressed genes in bulbils among different developmental stages (T1-T4). **(C)** Number of up- and down-regulated genes between bulbil developmental stage comparisions.

We further identified a total of 6,112 DEGs in a least one comparison (Supplementary Table S4, Fig. 2C). These results represented substantial differences in gene expression profiles during bulbil outgrowth. GO enrichment analysis was carried out to investigate their biological functions. We found that GO terms including response to abscisic acid, response to auxin, regulation of meristem growth, starch biosynthetic process, plasma membrane, cell wall and protein kinase activity, etc, were strikingly enriched (*P*<0.001) (Supplementary Table S5). Several significantly enriched (*P*<0.001) KEGG pathways for DEGs were suggested to be linked to starch and sucrose metabolism, plant hormone signal transduction, plant circadian rhythm (Supplementary Table S6, Supplementary Fig. S3). Accordingly, we then focused our interest on genes participating in these biological functions and metabolic processes.

### Light Stimuli and Circadian Clock During Bulbil Outgrowth

Several key photoreceptors perceiving the light signaling were here detected, including phytochrome A (phyA), phytochrome B (phyB) which presented lower expression levels in T1 and consistently higher levels in T4 (Fig. 3_I, Supplementary Table S7). For instance, three genes encoding phyB were up-regulated with an average of 3-fold (s) level in T4 as compared to T1. As a response to light stimulus, genes such as light-inducible protein (CPRF2) binding to G-box like motif (5’-ACGTGGC-3’), early light-induced protein (ELIP1) integrating the light-harvesting chlorophyll system, and light-dependent short hypocotyls (LSH4, LSH6) promoting cell growth, was activated in T1 and further elevated in subsequent developmental stages. Additionally, circadian clock regulators including CIRCADIAN CLOCK-ASSOCIATED 1 (CCA1) and LATE ELONGATED HYPOCOTYL (LHY) had higher expression levels in T1 as compared to other stages, suggesting that they can gate the rapidly altered light-quality responses.

**Fig. 3.**
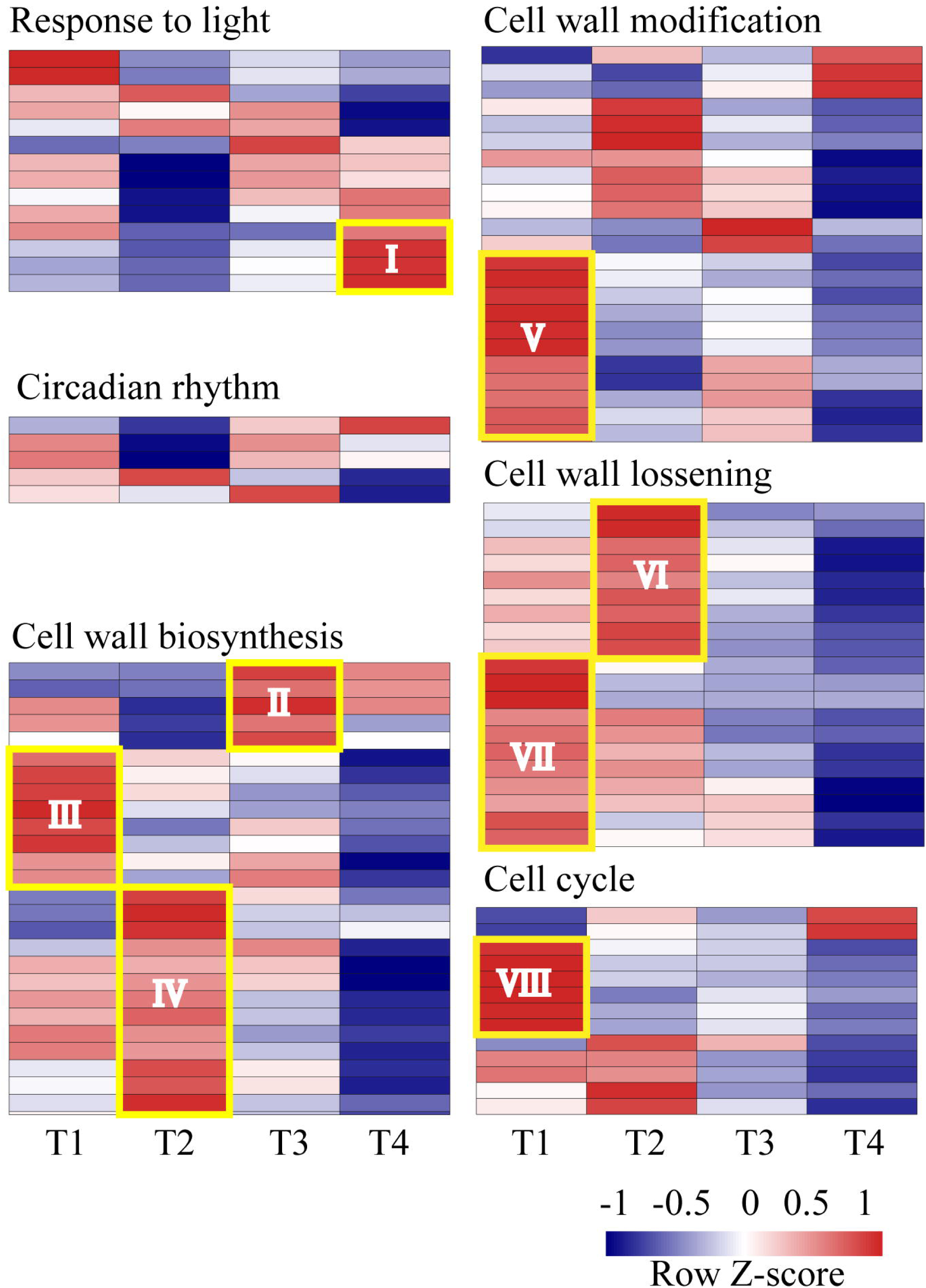
Expression profiles of DEGs involved in response to light, cell wall biosynthesis and modification, and cell cycle. The expression patteren of genes related to these biological functions was significantly distinct among different stages. Each row of the heat map represents an individual genes. The gene expression level is standardized into Z-score and colored in red and blue for high and low expression (see “Materials and Methods”).

### Differential expression of cell cycle genes during bulbil outgrowth

To better understand the control of cell cycle, we screened for DEGs involved in cell cycle and growth. We detected some marker genes for the control of cell division in bulbils, such as A-type, B-type and D-type cyclins (CYCA1, CYCA3, CYCD6), and cyclin-dependent kinases (CDKF1, CDKF4) (Fig. 3, Supplementary Table 8), which exhibited distinct expression changes. Most cyclins were up-regulated in the initial stage of bulbil formation (T1) (Fig. 3_VIII); in contrast, CDKF1 and CDKF4 showed down-regulated in T1 and further up-regulated in the later stage (T4). In addition, two homologous cyclin-dependent kinase inhibitor 4 (KRP4) were observed and presented relatively higher expression levels at T2 stage (Fig. 3), which can inhibit CDKs complex activity by phosphorylation and allow S phase progression (Verkest *et al*., 2005).

### Cell wall biosynthesis, modification during bulbil outgrowth

We totally identified 27 DEGs encoding biosynthetic enzymes for the building of primary and second cell wall (Supplementary Table S9). Of these genes, cellulose synthases or -like proteins (CESA7/8, CSLD2, CSLE6), and protein COBRA (COB, COBL7) (Fig. 3_III) showed higher expression levels in T1 compared to other stages. Galacturonosyltransferase or -like (GAUT3, GAUT7, GAUT14, GATL1, GATL3, etc) were expressed with higher levels in T2 (Fig. 3_IV). It was noted that multiple chitinases (CHI, CHI2, CHI5, etc) showed extremely high expression levels (> 7,067 FPKM). Unlike the greater part of genes in this cluster, chitinases exhibited higher expression levels in T3 (Fig. 3_II). We verified this expression profiles using qRT-PCR analysis (Supplementary Fig. S2).

Also, we detected 43 DEGs involved in molecular modification of wall network through degradation and loosening. For example, genes encoding endo-1,3(4)- b-glucanases (including EGase, MAN1, MAN9, glycosyl hydrolases family 17) and pectinesterases that regulate cell wall degradation had the highest expression levels in T1(Fig. 3_VII). Multiple xyloglucan endotransglucosylases (XTHs) showed relatively higher expression levels in the first two stages of bulbil formation (T1, T2) (Fig. 3_VI), which can loosen some wall-like networks. Besides XTHs, ten expansins (EXPs), as known cell wall loosening agents, were identified and most of them were up-regulated at the early stage of bulbil formation (Fig. 3_V).

### Starch and sucrose metabolic processes

KEGG enrichment analysis revealed that the pathway of starch and sucrose metabolism was the most enriched (Supplementary Table S6). We predicted some marker genes involved in starch synthesis, and identified genes encoding ADP glucose pyrophosphorylase (ADPG), small subunit of Glu-1-P adenylyltransferase (ADG), starch synthase (SS), glucan-branching enzyme (SBE), granule-bound starch synthase (GBSS). Some of these enzymes showed strong changes, and their abundances were commonly down-regulated in T1and sharply elevated in subsequent stages (Fig. 4). Particularly, SBE1, GBSS and ADG1 were strongly expressed. Additionally, we examined some genes involved in starch degradation, including genes encoding α/β-amylases (AMYs, BAMs), isoamylases (ISAs), phosphoglucan water dikinase (PWD), phosphoglucan phosphatase (PGP), beta-glucosidases (BGs) (Supplementary Table S10). Of particular interest genes AMY3, BAM9 showed highly expressed levels. Unlike most starch biosynthetic genes above, the two genes were up-regulated in T2 and successively down-regulated in T3 and T4.

**Fig. 4.**
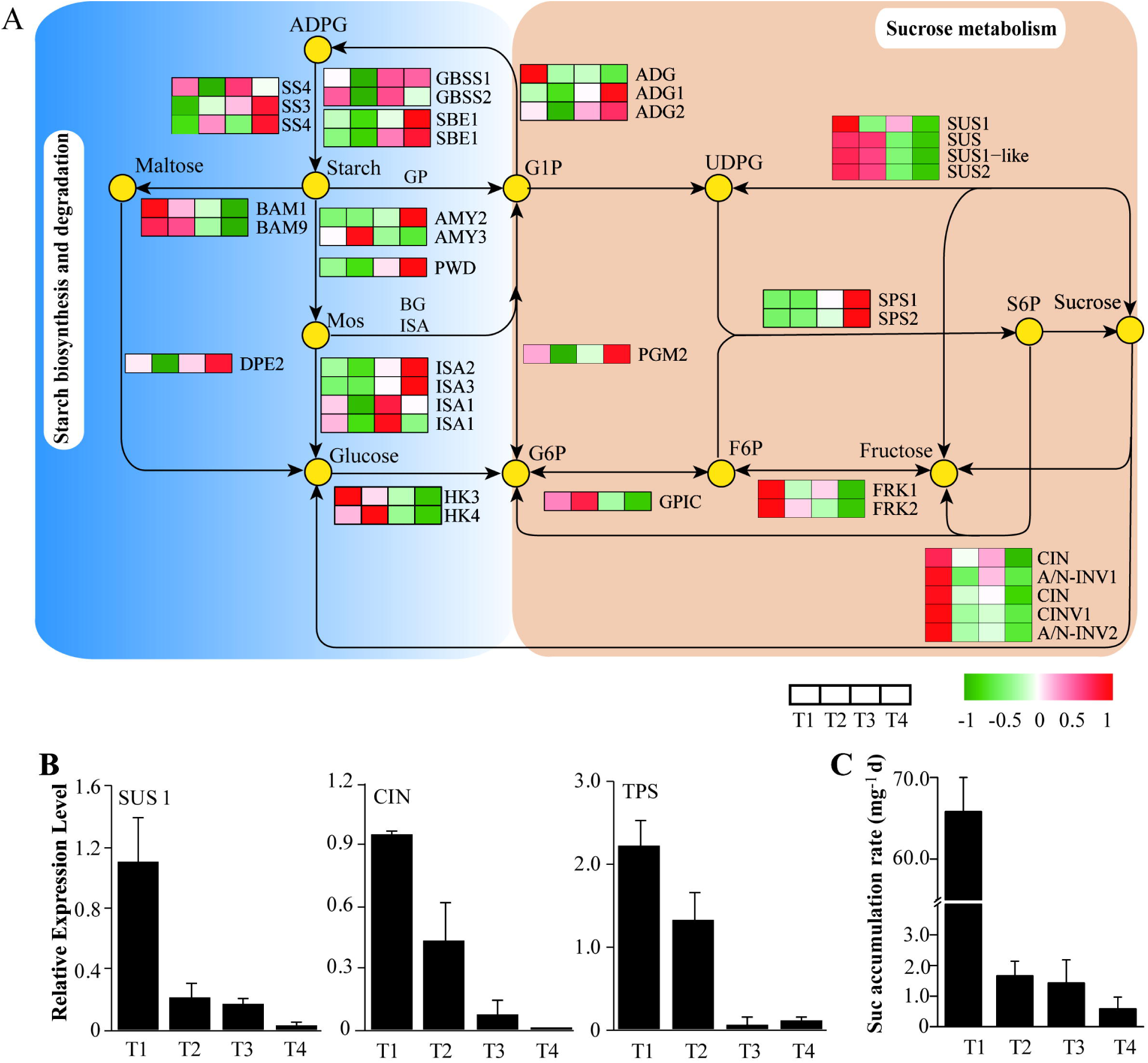
Sophisticated gene and metabolite regulation involved in carbohydrate pathways. **(A)** Schematic diagram of starch and sucrose (Suc) metabolsim, suggesting that the Suc biosynthsis pathway was highly up-rugulated at the early stage of bulbil outgrowth (T1). Heat maps next to the arrows represente the expression change of genes encoding corresponding enzymes for reactions. The expression level is standardized per gene into Z-score and colored in red and green for high and low expression. **(B)** Verfication of expression levels in genes encoding SUS1, CIN and TPS determined by qRT-PCR. **(C)** Sucrose accumulation in bulbils, demonstrating that sucrose is sharply required at the early stage of bulbil outgrowth (T1). Data show the mean±standard deviation (n=5). ADG, glucose-1-phosphate adenylyltransferase; ADPG, ADP glucose pyrophosphorylase; AMY, α-amylase; A/N-INV, neutral/alkaline invertase; BAM, β-amylases; BG, glucan endo-1,3-beta-glucosidase; CIN, cell wall invertase; CINV, cytosolic invertase; DPE, 4-alpha-glucanotransferase; FRK, fructokinase; F6P, fructose 6-phosphate; GBSS, granule-bound starch synthase; GPIC, glucose-6-phosphate isomerase; GP, glycogen phosphorylase; G1P, glucose-1-phosphate; HK, hexokinase; ISA, isoamylase; Mos, malto-oligosaccharides; PGM, phosphoglucomutase; PGP, phosphoglucan phosphatase; PWD, phosphoglucan water dikinase; SBE, glucan-branching enzyme; SPS, sucrose-phosphate synthase; S6P, sucrose 6-phosphate; SS, starch synthase; SUC, sucrose synthase; TPS, trehalose-phosphate synthase; UDPG, UDP-galactose.

Furthermore, there were multiple genes encoding Suc synthases (SUSs), Suc-phosphate synthases (SPSs), cell wall invertases (CINs) and alkaline/neutral invertases (A/N-INVs), representing key genes that participate in Suc synthesis and metabolism (Fig. 4A). The levels of transcripts encoding four SUSs were significantly up-regulated in T1 and gradually decreased in next stages. Particularly, SUS1(c130514.graph_c0) showed highly expressed level with the FPKM value of >2,000.

Meanwhile, there were several genes encoding hexokinases (HKs), fructokinases(FPKS), phosphoglucomutase (PGM), Glu-6-P isomerase (GPIC). Except PGM2 showing a greatly up-regulated expression level in T4, the genes of HK3/4, FPK1/2 and GPIC had a dramatic changes and exhibited higher expression levels in T1 stage. Of those genes associated with Suc metabolism, we detected three INV isoforms and two A/N-INVs that consistently presented up-regulated expression in T1. More importantly, we observed multiple members of trehalose-phosphate synthases (TPS) and trehalose-phosphate phosphatase (TPP) gene families involved in sugar signaling, their expression levels were successively up-regulated and down-regulated from T1 to T4 stage, respectively.

Given the importance of triggering axillary branch outgrowth by Suc, we further detected the expression levels of some key enzymes involved in Suc metabolism by qRT-PCR technology (Fig. 4B), and determined Sus accumulation in bulbils by detecting Suc contents (Fig. 4C). The genes SUS1, SUS1-like, and INV presented peak expression profiles in T1, which were consistent with that of transcriptomics analyses. We found that Suc was sharply accumulated in T1 and increased slowly in next stages.

### Dioscorin gene expression during bulbil outgrowth

We identified two members of dioscorin (Dio) family, DioA (c130377.graph_c3) and DioB (c135265.graph_c3) encoding diocorin protein. Phylogenetic analysis demonstrated that the two genes had high similarity with Dio sequences from other *Dioscorea* species (Supplementary Fig. S4) (Conlan, *et al*., 1998; Xue *et al*., 2012). It is noticeable that the two genes were expressed at fantastically high levels with >18,246 FPKM for DioA over bulbil outgrowth, and >9,878 FPKM for DioB. Both of them had increasingly expressed abundances during bulbil outgrowth. This expression changes were confirmed by qRT-PCR analysis (Supplementary Fig. S2).

### Transcriptional profiles of auxin synthesis, transport, and signaling components

To investigate the hormonal regulation during bulbil outgrowth, we analyzed the expression of genes involved in auxin biosynthesis, transport, and signaling (Fig. 5, Supplementary Table S11). We detected 12 auxin biosynthetic enzymes that were differentially expressed. Most of genes, including three TRYPTOPHAN SYNTHASE Alpha/ Beta (TSA, TSA-like, TSB1), two TRYPTOPHAN AMINOTRANSFERASE RELATED2/3 (TAR2, TAR3) and two FLAVIN-CONTAINING MONOOXYGENASES (YUCCA1), had higher expression levels in T1 relative to other stages (Fig. 5A). In addition, two INDOLE-3-ACETIC ACID-AMIDO SYNTHETASES (GH3.5, GH3.8) presented up-regulated expression in T1, which can catalyze the synthesis of IAA-amino acid conjugates (IAA-R) and provide a mechanism for depleting excess auxin in bulbils.

**Fig. 5.**
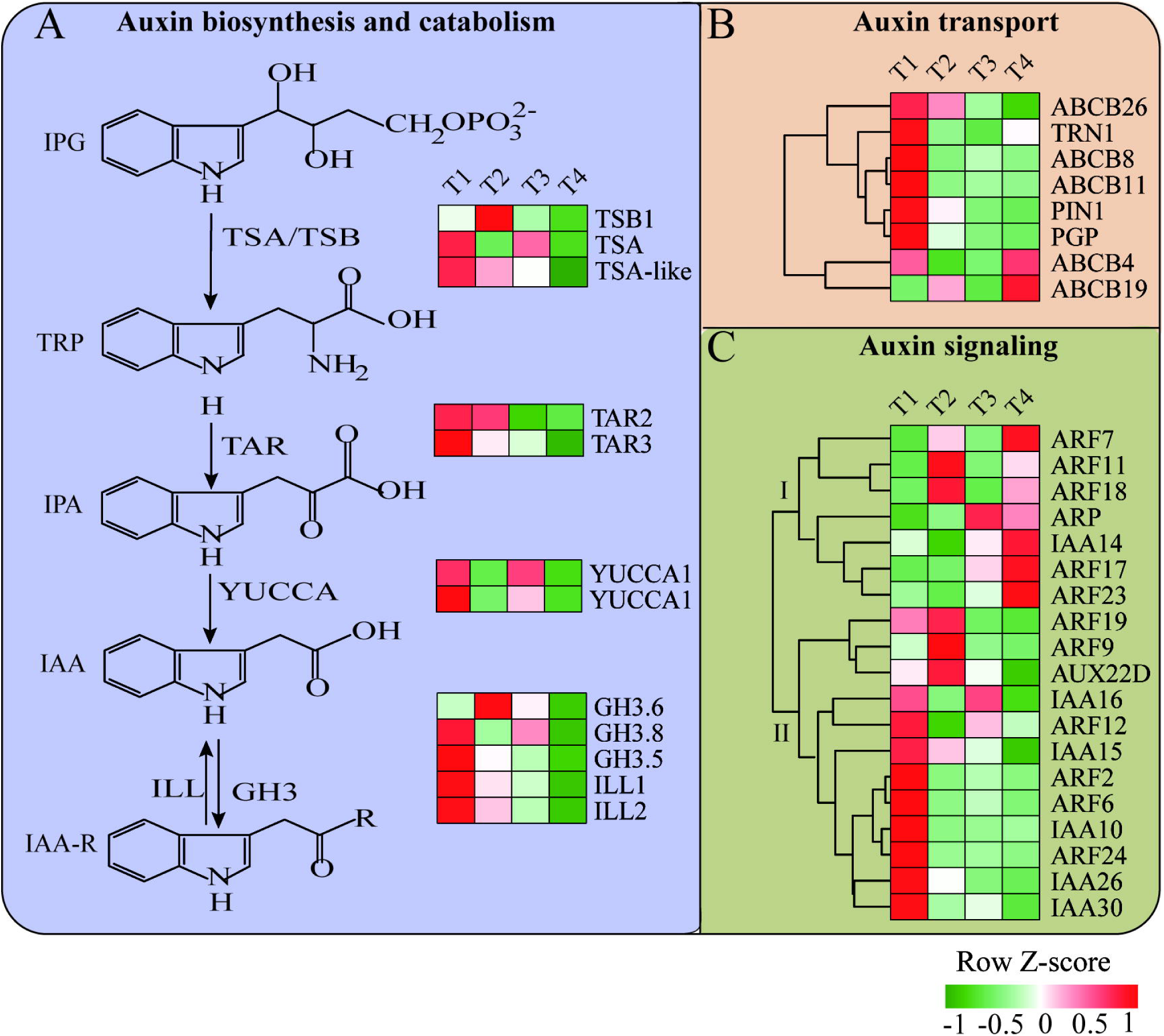
The regulatory framework of auxin-related genes. Hierachical clustering of expressed genes involved in auxin biosynthsis and catabolism **(A)**, transport **(B),** and signaling **(C)**, suggested that most genes were up-regulated at the early stage of bulbil outgrowth (T1). The expression level is standardized per gene into Z-score and colored in red and green for high and low expression.

Most auxin transporters showed increased expression levels in T1 relative to other stages (Fig. 5B). For instance, the expression levels of influx carrier protein TORNADO 1 (TRN1), transmembrane-targeted efflux carrier PIN1, other efflux carriers such as ATP-binding cassette subfamily (ABCB)-type transporters peaked in T1. In particular, ABCB19 (c133592.graph_c0) encoding P-glycoprotein (PGM) that regulates the polar auxin basipetal transport (Mravec *et al*., 2008), was highly expressed (154 FPKM) in T1 and sharply decreased to 54 FPKM in T2. Instead, only two ABCB transporters (ABCB4, ABCB19) were up-regulated in T4 at weak levels less than 15 FPKM.

Based on HCA, several key components of auxin signaling including AUX/IAA co-receptors, auxin-repressed protein (ARP), and ARF transcription factor binding with TGTCTC auxin response element (AuxRE), were grouped into two categories, and showed distinct expression patterns (Fig. 5C). Group I genes were more highly expressed in T4 relative to other stages. By contrast, group II displayed higher expression levels at the early stages (T1,T2). Among these genes, IAAs and AFBs genes were widely expressed. Specially, ARF6 and IAA15 exhibited relatively higher expression levels, which have been implicated in axillary shoot development in potato and tomato (Faivre-Rampant *et al*., 2004; Deng *et al*., 2012).

### Expression patterns of other plant hormone-related genes

We also conducted a profile analysis of genes that involved in other major hormones metabolism and signaling (Supplementary Table S11). We observed some genes encoding CK-deactivating gene, cytokinin oxidase 1/dehydrogenase (CKX1, CKX3, CKX9), and cytokinin ribosides (LOG1, LOG4) responsible for CK biosynthesis and catabolism. Most of them exhibited up-regulated expression levels in T1. In contrast, a gene encoding equilibrative nucleotide transporter 3 (ENT3) was highly expressed in T2 but was then strongly down-regulated in T4, which participates in CK transport. Multiple CK receptor genes encoding Histidine kinases (AHKs) and CK-inducible two-component response regulators (ARRs) were observed; they were up-regulated in T1 at moderate expression levels.

In addition, seven genes encoding zeaxanthin epoxidases (ZEPs) responsible for ABA biosynthesis were weakly expressed and showed down-regulated in T1 and then up-regulated in T4. Furthermore, we observed a set of ethylene-related genes, including biosynthetic genes encoding 1-aminocyclopropane-1-carboxylate oxidases (ACOs), receptor genes such as RTE1, EIN2 and EIN4, as well as transcriptional regulators (EFRs). Specially, genes EFR1, EFR2, EFR071 were highly expressed in T1 or T2 but were then strongly down-regulated in T4, demonstrating that the ethylene-inducible genes can be timely activated at the early stage of bulbil outgrowth. Similarly, several key genes in JA biosynthesis were also expressed with higher levels in the early stages (T1,T2) than in the later stages (T4). For example, FAD7, AOS1 and LOX6 that encode respectively omega-3 fatty acid desaturase, allene oxide synthase 1 and lipoxygenase, had the highest expression levels in T1. Additionally, a few DEGs involved in SA biosynthesis and signaling were identified and expressed at low levels, including genes encoding isochorismate synthase 2 (ICS2), SA-receptor proteins (SABP2, NPR1). both of them showed weak expression changes among different stages.

### Genes encoding transcription factors (TFs)

We observed 215 genes encoding for TFs that showed differentially expressed during bulbil outgrowth, and most of them were from AP2/ERF, WRKY, bHLH, NAC, MYB, MYB-Related, C2H2, C3H families (Supplementary Table 12). Given the fact that members in identical TF family may share similar functions, we calculated the expression proportion (P) of each TFs family relative to total expression level of all families over bulbil developmental stages (Supplementary Table 13). We found that AP2/ERF family showed highly expressed (P>5.7%), and families of C3H, WRKY, bZIP, NAC, bHLH and MYB, were moderately highly expressed (P>1.0%), suggesting that these families may play a more prominent role during bulbil outgrowth.

To display TFs expression profiles in detail, we clustered members of all TFs families into four distinct groups that represent four stage-specific expression patterns (Fig. 6). In cluster I, both of five members from C2H2 and C3H families exhibited specific expression and were strongly up-regulated in T4. In cluster II, 11 members of WRKY family represented a function category and showed dramatically increased expression in T2. Furthermore, AP2/ERF (14 members) and NAC (4 members) families were overrepresented in cluster III, both of which peaked in T3 stage. The enrichment of five members of MYB family was observed in cluster IV, these members showed strongly up-regulated in T1. To further explore the role of transcriptional regulators, we selected three TFs of MYB (C126446.graph-c0), WRKY (C125026.graph_c1) and NAC (C116834.graph_c0) due to their higher expression levels, and examined their expression patterns by in situ hybridization (Fig. 7). Consistent with the RNA-seq results, we found that all of them were specifically expressed in the AM initiation zone (a dome-shaped tissue) at T1 stage, and accumulated gradually decreased chromogenic signals in subsequent developmental stages, indicating key roles that play during bulbil initiation.

**Fig. 6.**
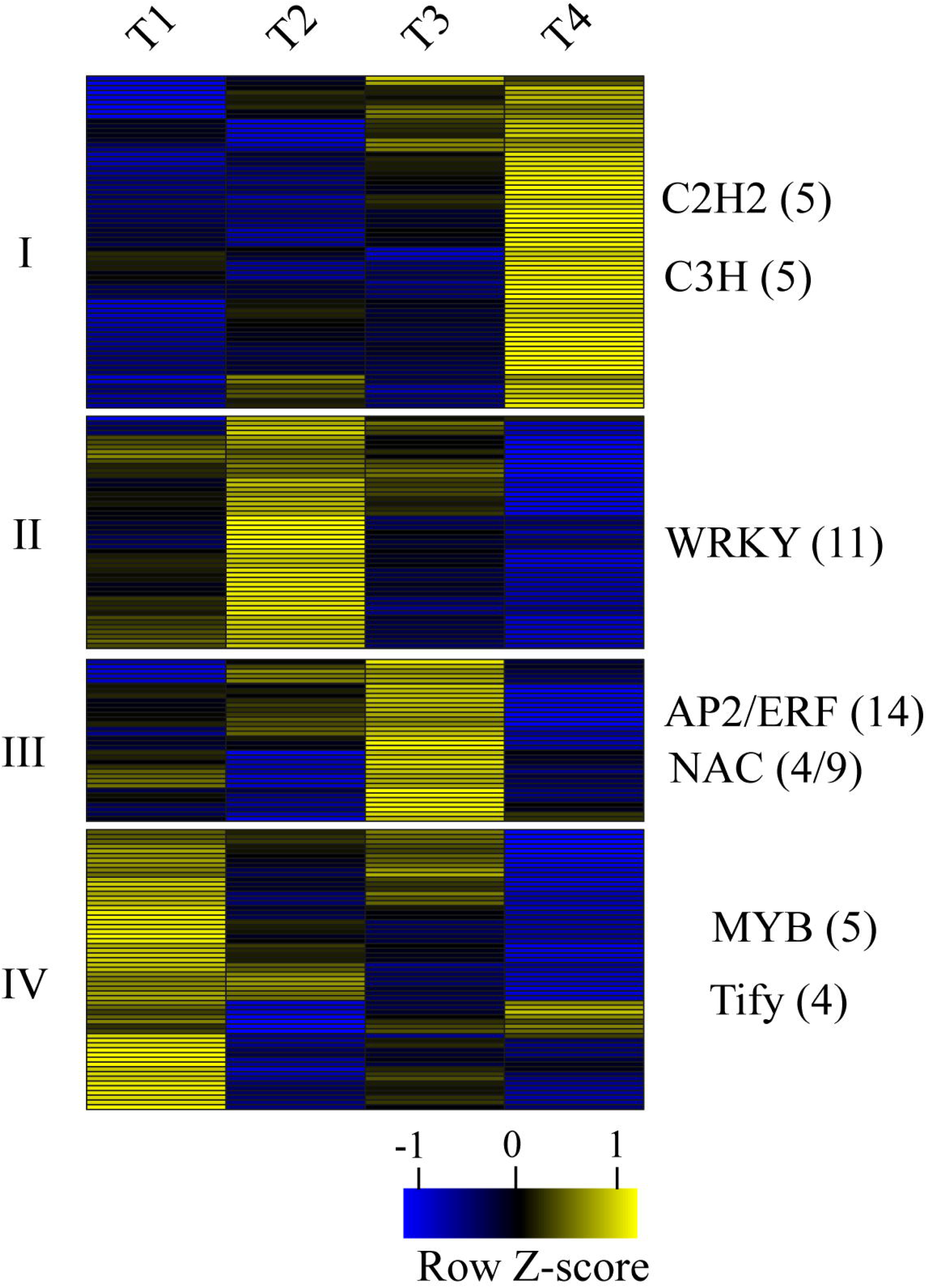
Expression profiles of transcription factors (TFs). Hierachical clustering seperated all differentially expressed TFs into four groups, indicating four stage-specific expression patterns. The TF families listed in the right of every group shows predominantly expressed in this group. The individual number represents the amount of member from TF families in the group. The gene expression level is standardized into Z-score and colored in yellow and blue for high and low expression.

**Fig. 7.**
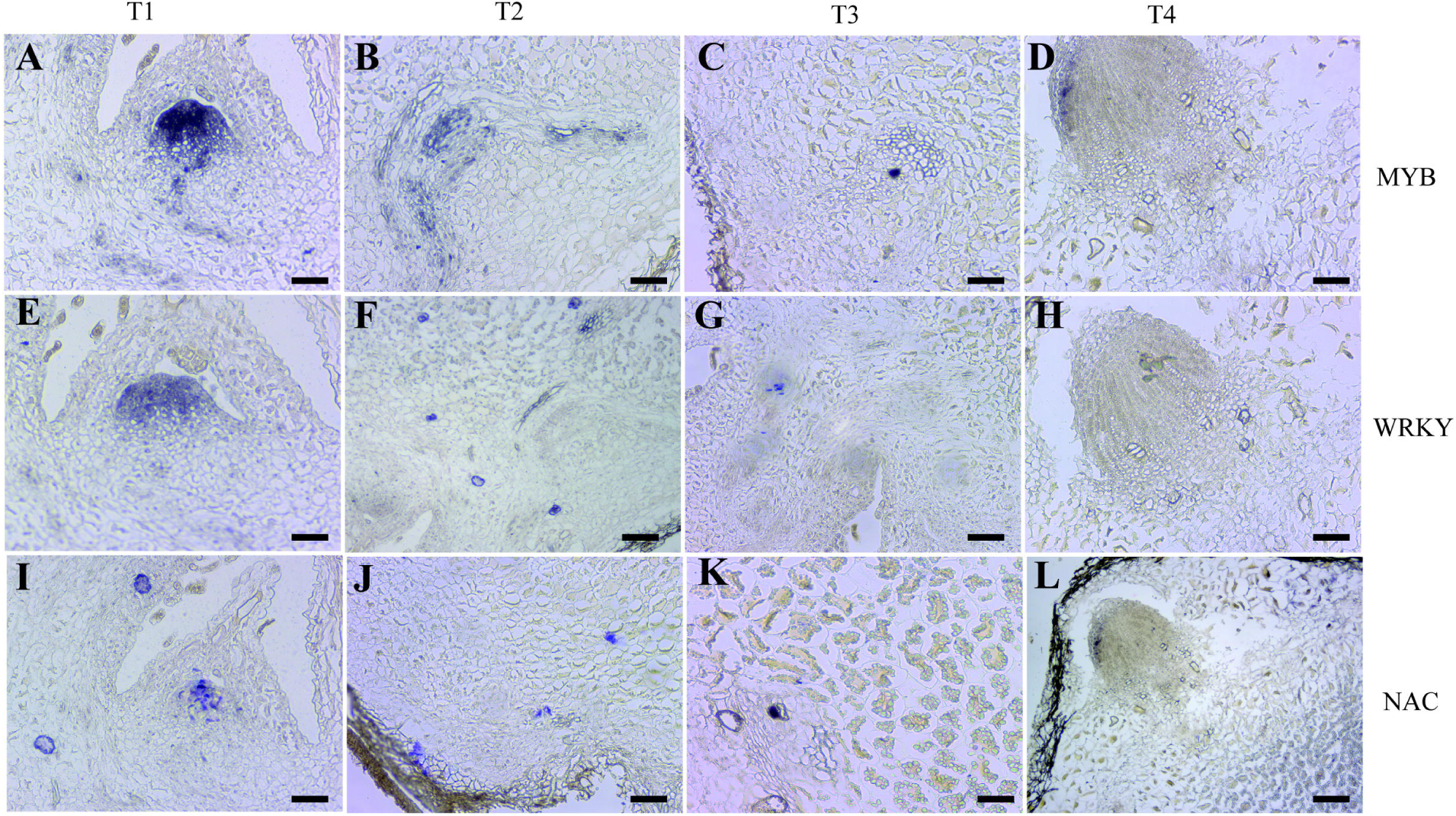
RNA in situ hybridization for MYB, WRKY and NAC transcription factors. Three transcription factors showed strongly accumulation in the AM initiation zone at the early stage of bulbil outgrowth (T1) (A, E, I). Bars=100 µm.

## Discussion

### Functional gene sets associated with yam bulbil outgrowth

Our transcriptome profiles confirmed that large sets of genes changed differentially between yam bulbil developmental processes, demonstrating these data can serve as resources to identify functional genes related to bulbil outgrowth. As shown in Figure 3_I, the gene cluster analysis showed the presence of a set of genes encoding photoreceptors including phy A, phy B and phototropins, which were strongly down-regulated in the young bulbil (T1). The function of the gene set (especially phyB) has been linked to a role of regulating the network of interacting light and hormones (Reddy, *et al*., 2013; Roman *et al*., 2016). Experiments in Arabidopsis has shown that phyB requires intact auxin signalling pathway to repress bud activity (Finlayson *et al*., 2010). Here, the outgrowth of young bulibils benefited from the inhibition of Phy B genes in the early stage. The result is in agreement with observations reported previously in sorghum plants (Kebrom *et al*., 2006; Kebrom and Mullet, 2016).

Another gene set is mainly represented by some cell cycle-related genes including CYCA1, CYCA3, CYCD6, CDKF1and CDKF4 (Supplementary Table S8). In the control of axillary buds, cell cycle-related genes performs a quality control function in promoting the resumption of meristem organogenic activity. The fact is directly supported by a variety of experiments that, increased gene expression of cell cycle in dormant buds from pea and Arabidopsis by decapitation stimulates bud outgrowth (Devitt and Stafstrom, 1995; Shimizu and Mori, 1998). Similar evidence has shown that the expression of several cell cycle genes (CYCB, CYCD2, CDKB) is decreased largely in sorghum buds by defoliation (Kebrom *et al*., 2010). In this study, a large set of cell cycle genes were strongly up-regulated in T1(Fig. 3VIII). This expression profile of genes increases the vitality of cell division in leaf primordia meristems and allows the resumption of organogenesis of young bulbils.

Meanwhile, several lines of evidences suggest that the expression of cell wall expansion related genes such as EXPANSINS, determines the rate of the initiation and elongation of premature axillary meristem and exerts a profound influence on plant development and morphology (Fleming *et al*., 1997). In a reported experiment from *Petunia hybrid* plant, over-expression of PHEXPA1 gene promotes axillary meristem release, whereas silencing PHEXPA1 produces opposite phenotypes (Zenoni *et al*., 2011). The enhanced expression of RhEXP by CK signal activates the SAM organogenic activity in rose axillary buds and further promotes the bud elongation (Roman *et al*., 2016). Consistent with these observations, we detected 11 gene set of EXPANSINS from EXPA and EXPB families, and found a distinctly up-regulated expression pattern in T1 (Fig. 3_V). Such result may be explained that the gene set of EXPANSINS contributes a unique role of maintaining cell enlargement during bulbil outgrowth and of building the special architecture form.

Starch constitutes most of the biomass of mature bulbil, accounting for 50-80% of its dry matter, and is a main trait being improved by breeding (Tamiru *et al*., 2008). As expected, we observed a set of ten marker genes involved in starch biosynthesis pathway (Supplementary Table S10, Fig. 4). Of particular interest is SBE and GBSS, which showed high expression levels in the later stages of bulbil formation (T3 and T4). This results are consistent with previous observation in maize embryo and endosperm development (Chen *et al*., 2014).

In addition to these conserved genes reported previously in axillary bud growth, we also focus genes that are highly expressed and specific to bulbil outgrowth. For example, two genes of Dio A and Dio B from Dio family encoding dioscorin protein showed successively increased expression profiles with extremely high levels (>9,878 FPKM) and, are homologous with some members cloned in other *Dioscorea* plant species (Xue *et al*., 2012). Dioscorin is the major storage protein and specific in tuber and bulbil from *Dioscorea* plants, which can support the new plant growth during germination or sprouting by supplying nitrogen (Xue *et al*., 2015). This finding strengthens the specialized role of Dio genes in bulbil outgrowth, implying that bulbil outgrowth can be somewhat understood to be the process of dioscorin protein accumulation.

### Bulbil outgrowth requires the coordinated control of Hormone-Related Genes

Understanding the expression of genes encoding integral components of hormone biosynthesis, metabolism or signaling facilitates efficient and directional links to hormone involvement during bulbil outgrowth. Here, the expression of genes encoding TSA/TAB, TAR and YUCCA enzymes involved in auxin biosynthesis were highly up-regulated in T1, suggesting that the produce of auxin is localized in bulbils in this stage. Consistent with our observation, localized auxin biosynthesis is required for axillary meristem initiation in maize and in Arabidopsis (Gallavotti *et al*., 2008; Zhao, 2010). Knockout of YUCCA genes leads to fewer branches due to the absence of axillary meristem (Cheng *et al*., 2006). On the other hand, ATC model supports that auxin need to be exported from axillary buds for its outgrowth by establishing the localized PAT steam in bud stem (not in main stem) (Blilou *et al*., 2005). The PAT steam is driven by PIN proteins belonging to a family of auxin efflux carriers that can facilitate auxin export out of cells (Petrášek and Friml, 2009). Arabidopsis mutants with more axillary growth increase PIN protein levels and the amount of auxin moving by PAT steam (Prusinkiewicz *et al*., 2009). In pea plant, increased auxin export from buds is accompanied by PIN1 polarization after decapitation, and further activates bud outgrowth (Balla *et al*., 2011). In accordance with these evidences, we observed the highly increased expression levels of multiple auxin efflux proteins including PIN and ABCB -type transporters in T1 stage (Fig. 4B), whereby these transport proteins facilitate transporting excess axuin and trigger bulbil outgrowth.

Outgrowth of axillary buds is positively correlated with CKs levels that are inhibited by the moving auxin in main stem (Ferguson and Beveridge, 2009; Domagalska and Leyser, 2011). The CK levels in chickpea buds increase 25-fold after decapitation, suggesting that CKs are necessary to initiate bud outgrowth (Turnbull *et al*., 1997). In some cases, CKs can stimulate bud outgrowth by a direct application to buds, even in the presence of apical auxin (Dun *et al*., 2012). In phy B sorghum mutants, reduced expression of genes involved in CK biosynthesis, and signaling leads to the resistance to bud outgrowth (Kebrom and Mullet, 2016). Mutations in rice CYTOKININ OXIDASE (CKX) that degrades CK, give rise to increased panicle branches and spikelet numbers in inflorescence (Ashikari *et al*., 2005). Conversely, mutations for CK-biosynthetic gene (LONELY GUY) in rice produce fewer branches (Kurakawa *et al*., 2007). In our study, we revealed the activation of CK synthesis genes (LOG1, LOG4) and the repression of degradation genes (CKX1, CKX3, CKX6, etc.) in bulbils at T1 stage (Supplementary Table S11). Also, an increased expression of CK transporter (ENT3) showed highly up-regulated in T1, which facilitates the CK flow from main stem to bulbils. Taken a whole, the expression of CK-related genes above could drive the promotion of bulbil outgrowth at the initiation stage.

ABA has also been thought to be a key component of regulating axillary organ growth. From a variety of plant species, the decline of ABA levels after decapitation precedes the onset of bud outgrowth (Zheng *et al*., 2015). In Arabidopsis, elevated ABA delays bud outgrowth and decreases elongating buds under low R:FR condition (Reddy *et al*., 2013; Yao and Scott, 2015). Several hypotheses has been postulated that ABA acts downstream of auxin and SL, possibly as a second messenger (Cline and Oh, 2006; López-Ráez *et al*., 2010). Independently of the presumption, it is undoubted that increased expression of genes involved in ABA biosynthesis is linked to repression of axillary bud growth. Consistent with previous reports, our data revealed that seven ABA synthesis genes (ZEP) are down-regulated in T1 and then strongly up-regulated in T4; and the degradation gene (CYP707A) exerted a reverse expression profile (Supplementary Table S11). Therefore, it is possible that these genes decreased ABA accumulation in bulbils at the initiation stage, thereby promoting bulbil outgrowth.

### Sucrose accumulation is the key for bulbil outgrowth

In addition to phytohormone signals, sugar (sucrose or analogues) as a novel player, contributes to the activation of axillary buds growth. In diverse plant species after decapitation, the progressive decrease of auxin levels in stem is too slowly to dominate the early bud formation, whereas sugars are speedily redistributed and enter buds to promote them growth (Barbier *et al*., 2015). Form a representative experiment in pea plant, the rate of polar auxin transport is 1cm·h^−1^ when loss of apical dominance, yet the outgrowth of bud reaches 40 cm at 2.5 h after decapitation (Mason *et al*., 2014). In our study, the rapidly accumulated rate of sucrose was observed in T1 (Fig. 4C), it is benefit to trigger the release of bulbils. Meanwhile, transcriptome analysis revealed that the expression of key genes (SUSs, CINs) involved in sucrose biosynthesis were highly up-regulated in the young bulbils (T1), which can timely unchoke the process of sucrose supply. More importantly, increased CIN expression can positively regulate axillary bud initiation by generating sugars for trophic uptake under interplay of light and phytohormones (Rabot *et al*., 2014).

On the other hand, sucrose functions as a critical signal through regulating the pathways involving T6P, HXK1 (O’Hara *et al*., 2013). Over-expressed HXK1 Arabidopsis lines show enhanced branching (Kelly *et al*., 2012). Particularly, the elevated T6P level has been implemented to be the signal that sugars increase influx into buds after decapitation, whereby buds from dormancy are released and elongated (Yadav *et al*., 2014; Figueroa and Lunn, 2016). T6P is synthesized by TPS and dephosphorylated to TPP. Over-expression of *E. coll* TPS (OtsA) in Arabidopsis results in the rise of T6P levels and further triggers the proliferation of shoot branching; instead, over-expression of *E. coll* TPP (OtsB) decreases both T6P levels and shoot branching (Schluepmann *et al*., 2003). Similar evidence demonstrated that constitutively affected Arabidopsis lines in synthesis and degradation of T6P, increases and reduces branching phenotypes, respectively (Yadav *et al*., 2014). In garden pea, T6P was attested as the signal of sucrose availability to promote outgrowth of axillary buds (Fichtner *et al*., 2017). In our study, we observed a successive up-regulated and down-regulated expression for TPS and TPP genes from T1 to T4 (Supplementary Table 11), which is more likely a consequence of enhanced sucrose signaling. These up-regulated TPS genes could increase T6P levels, thereby contributing to the promotion of bulbil outgrowth.

### Initiation of bulbil specifically expresses a large number of TF genes

The initiation of AMs is tightly linked to the activity of bud-specific regulators. Most of these genes belong to members of multiple TFs families, including R2R3 MYB proteins (REGULATOR OF AXILLARY MERISTERMS, RAX) from Arabidopsis, NAC domain TFs (CUP-SHAPED COTYLENONs, CUCs), a WRKY domain protein (EXCESSIVE BRANHES1, *EXB1*), GRAS domain proteins (LATERAL SUPPRESSOR, *LS*) in tomato, and TCP TFs (BRANCHED1/2, *BRC*1/2) in Arabidopsis and in sorghum (*TB1*) (Janssen *et al*., 2014; Yang and Jiao, 2016). Genetic studies from mutant plants have demonstrated that these transcriptional activators are specifically expressed in the boundary zone between leaf primordium and SAM, and control the fate of AMs initiation and the production of branches (Keller *et al*., 2006; Yang *et al*., 2012; Guo *et al*., 2015). For instance, loss of RAX1 gene encoding MYB37 in Arabidopsis, or its orthologous gene *BLIND* (*BL*) in tomato, fails to generate lateral buds during vegetative development, indicating the function of these genes being conserved (Keller *et al*., 2006; Naz *et al*., 2013). Similarly, WRKY TFs have been implicated in the control of axillary meristem (AM) initiation by transcriptionally regulating RAX genes. Over-expression of *EXB1* encoding WRKY71 increases excessive AM initiation and bud activities and, produces bushy and dwarf phenotypes (Guo *et al*., 2015).

In this study, we found that GO terms “ positive regulation of transcription “(*P*=8.6 E-12) and” regulation of meristem growth”(*P*=6.0 E-8) were significantly enriched in DEGs (Supplementary Table S5). We verified a large set of TFs genes over bulbil developmental stages, especially from members of AP2/ERF, WRKY, NAC, bHLH and MYB families (Fig. 6). Three representative TFs from WRKY, NAC and MYB were further confirmed to be highly expressed in the meristematic cell zone at the early stage of bulbil formation (Fig. 7). These data suggested that transcriptional regulators are required for the early step of bulbil expansion. However, the downstream genes that are controlled by them, still need to be explored by genetic approaches in future studies.

In conclusion, we have highlighted that long-distance signals (auxin, CK and ABA) play critical roles in controlling the bulbil formation from leaf primordium to outgrowth. Sucrose as a critical signal is strongly required at the early stage of bulbil formation through upregulating the T6P pathway. We have identified large sets of functional genes responsible for bulbil outgrowth, including genes related to light stimuli, cell division, cell wall modification and carbohydrate metabolism. Remarkably, some of these genes are novel and deserve special attention because of their extremely high expression levels, including Dio A/B proteins, starch synthetic enzymes and chitinases, which may be utilized to improve yam breeding program by genetic manipulation. Also, we have described the key role in triggering bulbil initiation regulated by transcriptional regulators. Overall, our work presented here allows, to our knowledge for the first time, a full overview of transcriptomic profiling for yam bulbil outgrowth, and provides key genetic factors underlying bulbil outgrowth regulation.

## Acknowledgements

This study was supported by Zhejiang Provincial Key Laboratory for Genetic Improvement and Quality Control of Medicinal Plants (No. 2011E10015) in China, and Zhejiang Provincial Applied Research Program for Commonweal Technology (No. 2016C32SA300040).

## Author contributions

ZG.W conceived the program and designed the experiment, and supervised the writing. W. J analyzed the transcriptome data and coped with the figures. ZM. T, XZ. G and WH. Y performed most of the experiments and wrote the manuscript with their contributions. XJ. P helped to analyze bioinformatics. All authors read and approved the final manuscript.

## Supplementary data

Supplementary data are available at *JXB* online.

**Fig. S1.** Pearson correlation relationship between biological replicates.

**Fig. S2.** Validations of gene expression profiles by qRT-PCR.

**Fig. S3.** Enriched KEGG pathways.

**Fig. S4.** Phylogenetic analysis for Dio A and Dio B genes.

**Fig. S5.** Expression dynamics of TF families.

**Table S1.** List of the primer sequences used in this study.

**Table S2.** Summary of RNA-seq reads in this study.

**Table S3.** Assembly statistics for the bulbil transcriptome.

**Table S4.** Lists of differentially expressed genes.

**Table S5.** List of significantly enriched top GO terms.

**Table S6.** Lists of significantly enriched KEGG pathways.

**Table S7.** Expression of genes related to light stimuli and circadian clock.

**Table S8.** Expression of genes related to cell cycle.

**Table S9.** Expression of genes related to cell wall biosynthesis, modification and loosening.

**Table S10.** Expression of genes related to carbohydrate metabolsim and sucrose signaling.

**Table S11.** Expression of genes related to hormone metabolism, transport and signaling.

**Table S12.** Lists of differentially expressed TFs.

**Table S13.** The expression proportion of the identified TF families.

